# GraphPart: Homology partitioning for biological sequence analysis

**DOI:** 10.1101/2023.04.14.536886

**Authors:** Felix Teufel, Magnús Halldór Gíslason, José Juan Almagro Armenteros, Alexander Rosenberg Johansen, Ole Winther, Henrik Nielsen

## Abstract

When splitting biological sequence data for the development and testing of predictive models, it is necessary to avoid too closely related pairs of sequences ending up in different partitions. If this is ignored, performance estimates of prediction methods will tend to be exaggerated. Several algorithms have been proposed for homology reduction, where sequences are removed until no too closely related pairs remain. We present GraphPart, an algorithm for homology partitioning, where as many sequences as possible are kept in the dataset, but partitions are defined such that closely related sequences always end up in the same partition. Evaluation of GraphPart on Protein, DNA and RNA datasets shows that it is capable of retaining a larger number of sequences per dataset, while providing homology separation quality on par with reduction approaches.

## Introduction

When constructing a statistical or machine learning (ML) method, it is crucial to test its predictive performance on a test set - a dataset that was not involved in calculating the parameters during the training process. This is necessary to measure whether the method can *generalize* from the examples in the training set and produce useful output for previously unseen data. If a model reproduces its training examples in too much detail, it uses its parameters to fit not only the common pattern in the data, but also the individual noise in each data point. When this happens, the performance on the test set goes down, and the model is said to be *overfitted* (*1*). The common approach to monitor potential overfitting is to split the data into separate sets that are used for training and testing. If the model has *hyperparameters* such as architecture, learning rate, and length of the training process, three datasets should be used: a training set used for optimizing the model parameters, a validation set for optimizing the model architecture and training process, and a test set for measuring the performance (*2, 3*).

An extension of this approach to obtain more reliable estimates is *k*-fold cross-validation, where the data is split into *k* partitions so that the training procedure can be repeated multiple times with the training, validation and test sets swapped (*3*). If all combinations of training and validation set are tried for each test set so that *k* × (*k*-1) training runs are performed, it is termed *nested* cross-validation (*4*).

This problem is not unique to computational biology. In fact, all ML applications rely on estimating performance on test data that was not used for training or optimization. Typically, this can be achieved by splitting the data randomly. However, for biological sequences this approach is severely flawed: Sequences are evolutionarily related, and random splitting would lead to close homologs of training sequences being present in the validation and test data. This should be taken into account either by reducing the dataset until no pairs of too closely related sequences remain before splitting it into folds (*homology reduction*) or by ensuring that no too closely related pair of sequences end up in different folds (*homology partitioning*). This has been recognized quite early in the history of bioinformatics (*5, 6*).

However, it is non-trivial to define when sequences are neighbors or “too closely related”. This question has two parts: Which measure of similarity should be used, and what is an acceptable cutoff for this measure of similarity? The standard way of comparing two sequences is to do a pairwise alignment, either global (*7*) or local (*8*). Then, the percentage of identical nucleotides or amino acids can be calculated, and this percentage is often used as a measure. However, if local alignment is used, it is necessary to take into account the alignment (overlap) length as well. An 80% identity is not significant if it is just four out of five positions, but if it is 40 out of 50, it is highly significant.

One approach to a non-arbitrary definition is to identify a cutoff above which the problem could be better solved by alignment than by machine learning. In a 1991 paper, Sander and Schneider defined that this was the case for protein structure prediction, if more than 70% of the residues in a local alignment had the same secondary structure assignments. This was used to define a length-dependent percent identity cutoff which was *t*(*L*) = 290.15*L*^-0.562^ if *L* was between 10 and 80, and 24.8 if *L* was larger than 80. Alignments shorter than 10 were never considered neighbors (*5*).

Lund and coworkers revisited this definition in 1997, but found the formula for the threshold to be 290*L*^-0.5^ for all values of *L*. However, they observed that an almost identical performance was obtained using a constant alignment score as cutoff. They additionally aligned versions of the sequences where the amino acids had been shuffled in random order and showed that the percent identity of these alignments were distributed up to this length-dependent cutoff, very rarely surpassing it (*9*).

A similar analysis on a different prediction task was carried out by Nielsen and coworkers in 1996. They made pairwise alignments of N-terminal fragments of proteins containing signal peptides and considered whether the signal peptidase cleavage sites were precisely aligned. In contrast to the aforementioned works, they chose to define the cutoff as a certain absolute number of identical amino acids (*10*).

A different approach is to use the fact that local alignment scores for unrelated sequences follow an extreme value distribution (*11*). If the cumulative distribution of scores is transformed by a double-logarithmic function: ln(-ln(1-P_score>*x*_)) and plotted against *x*, it will show a straight line for unrelated sequences, and the cutoff can then be placed where the curve deviates significantly from the straight line (*12*).

Alignment scores are very dependent on the scoring system (substitution matrix and gap penalties) (*11*) and are not very intuitively understandable. In practice, the threshold is therefore most often expressed as a percent identity. When using local alignment, this should be evaluated together with alignment (overlap) length as described above. Instead, the number is often calculated as the number of identical letters in the overlap divided by the length of the shorter sequence (instead of by the length of the overlap).

Two widely used algorithms for homology reduction were published by Hobohm and coworkers in 1992 (*6*). Algorithm 1 (“select until done”) starts with a list of sequences (which may be ranked according to some quality criterion) and works by selecting the first sequence and removing all its neighbors, then selecting the next sequence from the list and so on, until the list is exhausted. Algorithm 2 (“remove until done”) works by removing the sequence with the largest number of neighbors first, then recalculating the number of neighbors for each sequence, and continuing to remove the sequence with the largest number of neighbors until no pair of neighbors remain. Algorithm 2 typically preserves more sequences in the final set, but requires calculating all pairwise similarities, while Algorithm 1 only requires a subset of the distances.

An alternative to Hobohm’s two algorithms is the “greedy-min” algorithm which selects the sequence with the smallest number of neighbors, removes all its neighbors, recalculates the number of neighbors for each sequence, and again selects the sequence with the smallest number of neighbors and repeats until all sequences have either been selected or removed (*13*). It tends to preserve more sequences than Hobohm’s Algorithm 2, but in practical tests, we find that the difference is negligible (results not shown).

As mentioned, a drawback to the Hobohm 2 algorithm is the need to calculate all pairwise distances or similarities. To deal with this, a number of faster approximative clustering algorithms have been introduced, notably CD-HIT (*14, 15*), HHblits (*16*), and MMseqs (*17, 18*). A common feature of these algorithms is that they designate a *representative* sequence of each cluster and assign a cluster identifier to each sequence. It is important to stress that these algorithms can be used for homology reduction by selecting only the representative sequences, but *not* for homology partitioning, since the maximum similarity criterion is only applied to the representative sequences, not to the other members of the clusters.

Homology partitioning algorithms are largely missing from the literature. One exception is the recent article by Petti and Eddy about constructing benchmark sets using graph algorithms (*19*). However, that work focuses on benchmarking remote homology detection methods, so the objective there is to split each *family* of proteins into dissimilar training and test sets. In contrast, our objective is to split an entire dataset into dissimilar subsets.

We define the homology partitioning problem as follows: Given

- a set of sequences with associated class labels,
- a maximum similarity threshold and
- the number of partitions to generate,

find an assignment of sequences to partitions so that

I. no sequence in a partition has identity above the threshold to any sequence in any other partition,
II. all partitions have approximately the same number of sequences for each label and
III. as many sequences as possible are retained in the partitioned dataset overall.

To address this problem, we developed GraphPart, a graph clustering algorithm that finds the partitions by first finding and merging clusters, followed by an iterative reassignment and removal step. It consists of the following four steps (Figure 1):

**Figure 1.**
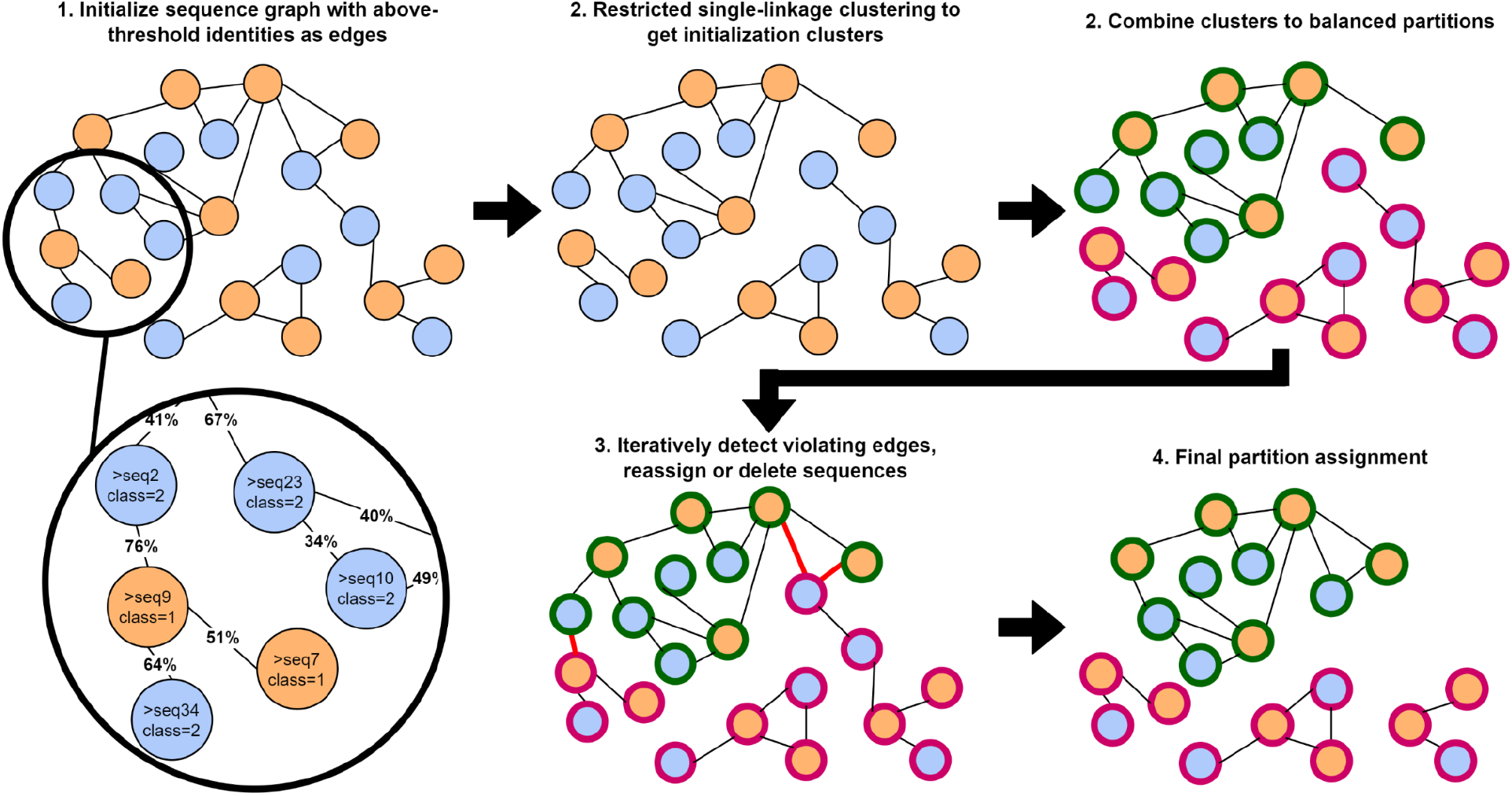
The GraphPart algorithm. Given a graph of sequences of two classes (orange and blue), with edges between sequences indicating identity above a chosen threshold, restricted single-linkage clustering is applied to yield class-balanced clusters. These clusters are combined to yield balanced partitions (green and red) of the required sizes. Given the partitions and the original sequence graph, sequences are iteratively removed or moved between partitions until there are no edges connecting the partitions, eliminating homology between different partitions.

I. A graph of all sequences is constructed by computing all pairwise alignments using a suitable alignment program, with the resulting percent sequence identities (homology) becoming the graph’s edge weights.
II. Clusters of homologous sequences are obtained by applying a restricted version of agglomerative single-linkage clustering that
  A. uses a predefined similarity threshold,
  B. will not merge clusters if the resulting cluster becomes larger than the expected size of a partition and
  C. will not merge clusters if the resulting count of any class becomes larger than its expected count per partition.
III. The clusters are iteratively assigned to the partitions such that the number of sequences of any label is approximately the same across partitions.
IV. Finally, a heuristic is applied to iteratively reassign or remove sequences until no sequences belonging to different partitions have an identity above the threshold.

As the similarity measure, we have chosen to use *global alignment* percent identity. As mentioned, the measure of choice is often percent identity from a local alignment, where the denominator is not the alignment (overlap) length, but the length of the shorter of the two sequences (e.g. in CD-HIT). A global identity measure will be smaller than or equal to this measure, because the global alignment length is at least as long as the longer of the two sequences. Therefore, using global percent identity allows for unbiased evaluation of algorithms on equal terms.

## Materials and Methods

### Datasets

To evaluate the performance of GraphPart in representative applications, we gathered multiple datasets of DNA, RNA and protein sequences (Table 1). All datasets were used in previously published machine learning predictors. For datasets that are already homology reduced in their published versions, we recreate the original datasets according to the instructions laid out in the respective paper. For DNA, we choose datasets of human TATA promoter sequences used for the training of DeePromoter (*22*) and of DNA regions binding to histone H3 (*23, 24*). For RNA, we recreate the non-coding RNA classification dataset used in nRC (*25*) by extracting sequences from Rfam (*26*). As a second dataset, we recreate a dataset used for RNA backbone angle prediction in SPOT-RNA-1D (*27*) by querying the Protein Data Bank (*28*).

**Table 1.** Summary statistics of the datasets used for benchmarking.

| Dataset | Type | Size | Maximum length | Classes | Threshold |
| --- | --- | --- | --- | --- | --- |
| DeePromoter | DNA | 2,929 | 300 | 1 | 80% |
| Histone | DNA | 14,965 | 500 | 2 | 80% |
| nRC | RNA | 6,500 | 758 | 13 | 80% |
| SPOT-RNA-1D | RNA | 1,227 | 2,880 | 1 | 80% |
| NetGPI | Protein (truncated) | 3,618 | 100 | 2 | 30% |
| DeepLoc | Protein | 14,004 | 13,100 | 11 | 30% |
| SignalP 5.0 | Protein (truncated) | 20,758 | 70 | 4 | 30% |
| NetSurfP 2.0 | Protein | 30,046 | 2,128 | 1 | 25% |

For proteins, we investigate performance separately for truncated protein sequences and full proteins. Truncation is often applied when it is already known a priori in which region of the protein the feature of interest is present. We use signal peptides (70 N-terminal amino acids) from SignalP 5.0 (*29*) and GPI anchors (100 C-terminal amino acids) from NetGPI (*30*). For full proteins, we use the eukaryotic subcellular localization dataset from DeepLoc (*31*) and the secondary structure datasets from NetSurfP 2.0 (*32*), which we recreate in its non-reduced state by obtaining Protein Data Bank data at 95% identity from PISCES (*33*). We partition protein sequences at 30% and RNA sequences at 80% identity. For NetSurfP, as it is a structural prediction task, we use a reduced threshold of 25%. For DNA, there is no commonly applied percent identity threshold to reduce homology. As 80% is the minimum achievable threshold of CD-HIT-EST, we also use 80% for DNA.

### Baselines

For each dataset, we compare the performance of GraphPart to the commonly used programs CD-HIT (*14*) and MMseqs2 (*20*). We benchmark two different application modes: *homology reduction* and *homology partitioning*. We cluster all sequences to the given identity threshold according to the programs’ tutorials. For protein data with CD-HIT, we first cluster with cd-hit at 90% identity, then at 60% identity, followed by psi-cd-hit to the final threshold. For nucleotides, we cluster with cd-hit-est directly at the final threshold. In all cases, -g is set to 1 for slower, more accurate clustering. For MMseqs2, we run mmseqs easy-cluster to cluster directly at the given identity threshold. MMseqs2 supports different clustering modes. In our initial experiments we found no clear effect of the clustering mode on separation quality, so we use the default mode 0 in all experiments.

In the homology reduction mode, we only retrieve the representative sequence for each cluster and partition this reduced set of sequences using the StratifiedKFold method included in Scikit-learn (*34*). For homology partitioning, we assign all sequences in a cluster to a partition using a heuristic assignment algorithm. We iterate over all clusters, assigning each cluster to a partition so that label ratios and partition sizes are as balanced as possible. We note that the latter approach is not expected to result in strict homology separation, but is still capable of reducing similarity between partitions compared to random split strategies.

### GraphPart Algorithm

The four steps in the GraphPart algorithm are:

I. GraphPart needs to compute all pairwise sequence identities in the input dataset *S*. For exact Needleman-Wunsch alignments, we use needleall from the EMBOSS (*21*) package with the respective defaults for protein and nucleotide sequences. When using MMseqs2 (*20*) alignments, we compute all-vs-all alignments without a prefiltering step as documented in the MMseqs2 user guide. We return all alignments that exceed the given identity threshold, without filtering for e-values. As the default denominator for sequence identity (length of the local alignment including gaps) can give misleading results when not using coverage controls, we use the length of the shorter sequence as the denominator. When aligning nucleotide sequences, the prefiltering step cannot be omitted and is thus executed with the sensitivity set to maximum.

The GraphPart partitioning algorithm operates on real-valued distance metrics. Sequence identities ranging from 0 to 1 are converted to distances as d(a,b) = 1-identity(a,b). The partitioning threshold undergoes the same conversion. Alternatively, GraphPart can accept any precomputed distance or similarity metric and skip the alignment step.

The result of this step is a graph with sequences *S* as nodes and distances as edge weights *D*.

II.The sequences are then clustered using a restricted form of single linkage clustering. Each sequence entity in *S* is initially regarded as a separate, or singleton, cluster. *D* is sorted in ascending order and iterated over to connect pairs of clusters, forming larger clusters. If a connection between two clusters would result in a cluster whose size exceeds the expected partition size 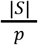 where p is the number of partitions, the clusters are not combined. Likewise, if the connection would result in a cluster having more sequences of any label *l* ∈ *L* than the expected value 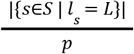, the clusters are not combined. If a pairwise distance exceeds the threshold, the procedure is stopped.

III.The clusters generated in the previous step are partitioned for balanced label composition. The desired number of partitions are initialized as empty sets. The list of clusters is iterated over, and all sequences of a cluster are assigned to one of the partitions. The partition to be assigned to is chosen so that the imbalance in the number of sequences of a label *l* across partitions is minimized after assignment.

IV.The last step is the removal phase, for which pseudocode is given in Supplementary Code 1. Sequences that have a distance lower than the threshold to any sequence in another partition are iteratively removed from *S* together with the corresponding pairwise distances being removed from *D*.

To achieve this, the following procedure is repeated starting from iteration *i* = 1: For each sequence *s* ∈ *S*, the number of connections below the threshold to its own partition and to the other partitions is computed. If *s* has more connections to sequences in another partition than in its own, *s* is moved to that partition. While iterating over *S*, we compute *c*, the total number of sequences having cross-partition connections below the threshold. After processing all *s* ∈ *S*, the c * log10(i/100)+1sequences with most cross-partition connections below the threshold are removed from *S*. The multiplier with which we multiply *c* is a heuristic scaling factor that ensures that removal proceeds less aggressively in early iterations. The procedure is repeated as long as c>0, incrementing *i* with +1after each iteration.

GraphPart also accepts binary priority labels to indicate which sequences should be removed first, if removal is required. When using such labels, the removal procedure is first performed in a restrained mode, only removing low-priority sequences until there are no more cross-partition connections for this group. If there are still cross-partition connections remaining, the removal procedure is then repeated in the unrestrained mode.

In addition to finding balanced k-fold partition assignments, GraphPart is also capable of generating train-validation-test splits, e.g. reserving 10% of the data for testing and 5% for model validation. To achieve this, we first partition the data into *n* partitions, with *n* being a number such that it yields natural numbers when multiplied with both the test and the validation ratios. We select the maximally connected *k* of the *n* partitions to yield the train set, with k=n* (100% – test % – validation %). Maximum connectivity is computed by counting all cross-partition edges exceeding the threshold. The same is then done with the remaining unassigned partitions to yield the test set. The final remainder becomes the validation set. After reassignment of all sequences according to their new split memberships, we proceed with the removal phase as in the standard case.

### Evaluation

For evaluating homology separation quality, we compute the maximum cross-partition pairwise identity for each sequence. Successful separation implies that there is no sequence in the dataset that has a cross-partition pairwise identity higher than the defined threshold.

We use EMBOSS needleall (*21*) for pairwise Needleman-Wunsch global alignments. Given a partition assignment, we align each sequence in a partition to all sequences in any other partition to find the sequence’s maximum cross-partition pairwise sequence identity. We leave all alignment parameters at their defaults, using EBLOSUM62 for protein and EDNAFULL for nucleotide alignments.

To evaluate runtime performance, GraphPart was executed on a 12-core Intel Xeon Gold 6126 HPC node, with the full node reserved for GraphPart execution for comparability. Execution times were logged directly by the GraphPart Python script. For both the sequence length and dataset size runtime evaluations, we upsampled the NetGPI dataset. For size, we sampled sequences with replacement to the desired total number. For length, we first trimmed the sequences to 50 amino acids and repeated the trimmed sequence N times to yield sequences of the desired length.

## Results

Throughout all investigated datasets, we find that performing homology partitioning with full clusters fails to achieve homology separation (Figure 2). This result is expected, as the procedure essentially only maximizes similarity within clusters in a partition, rather than minimizing similarity between partitions. However, we also find that homology reduction, which is expected to minimize similarity, can leave a significant amount of cross-partition NW identities above the threshold. In all datasets, reduction by CD-HIT yields better results than MMseqs2, but still falls short of achieving complete separation. Moreover, we observe a dependency on sequence length: Both MMseqs2 and CD-HIT see a significant decrease in separation quality when using shorter, truncated sequences instead of full proteins (Figure 2A, B). As expected, GraphPart achieves perfect separation irrespective of sequence length when using NW alignments. Additionally, using GraphPart with MMseqs2 alignments results in better separation than directly applying MMseqs2 or CD-HIT for homology reduction.

**Figure 2.**
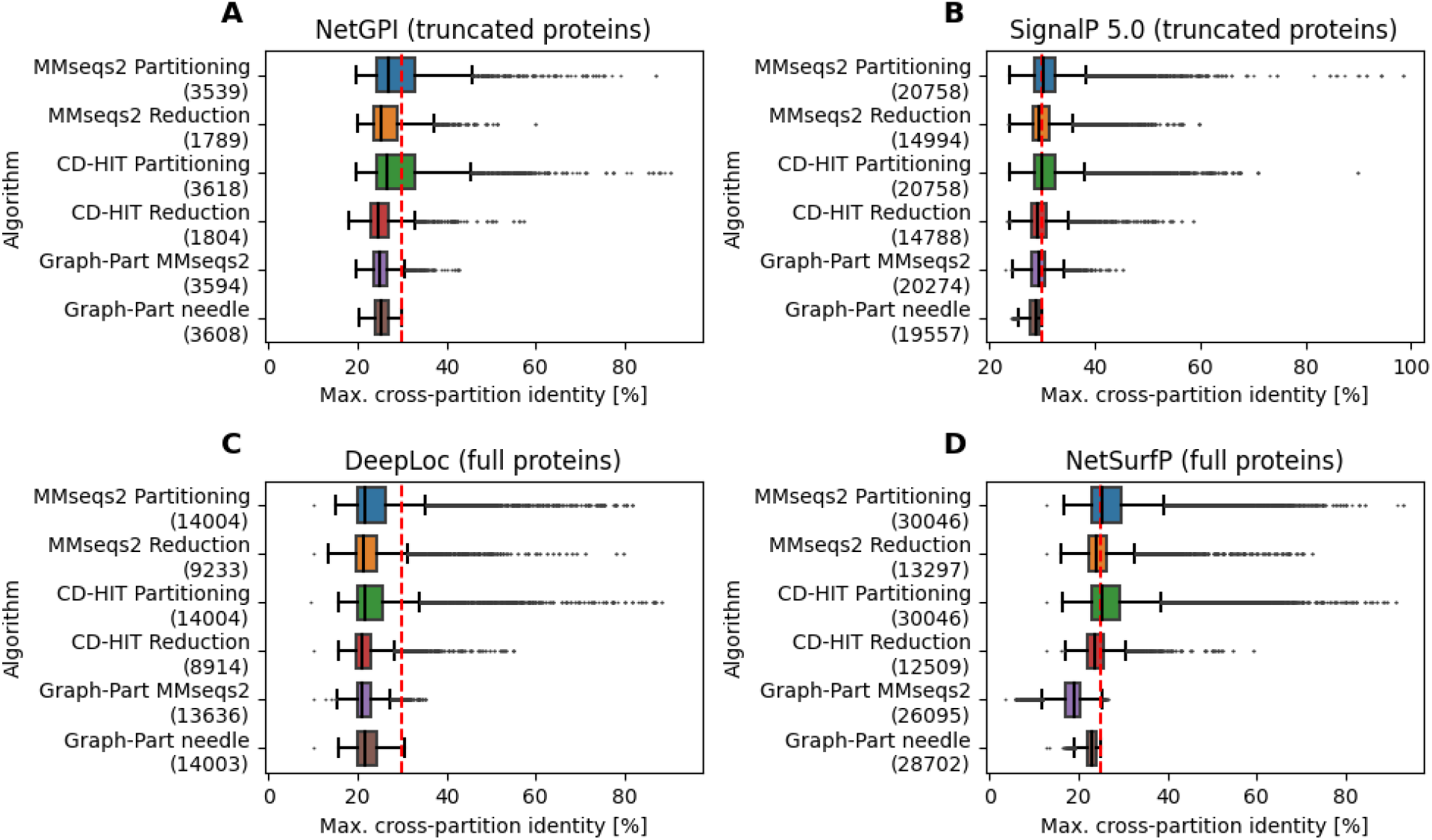
Benchmarking results on protein datasets. Datasets were partitioned into five folds. The target pairwise identity threshold is indicated as the red dashed line. All algorithms were run for the generation of five equal-sized balanced partitions.

Besides achieving better separation than homology reduction approaches, GraphPart is capable of retaining up to 50% more sequences in its partitions. This is due to GraphPart only removing sequences until there are no more pairwise identities between different partitions, rather than removing sequences until the pairwise threshold is obeyed by all remaining sequences in homology reduction. We further investigate the retention capability by benchmarking against Hobohm’s algorithm 2 for homology reduction with NW alignments (*6*). While Hobohm’s algorithm continuously reduces the number of sequences with decreasing identity thresholds, GraphPart retains most sequences until a critical threshold is reached at which large-scale removal becomes unavoidable (Figure 4). In practice, we find that this critical threshold lies lower than commonly used homology cutoffs.

When benchmarking on nucleotide data, we observe that MMseqs2 homology reduction can fail to remove identities above the threshold (Figure 3). Overall, RNA datasets show a higher redundancy than our protein datasets, with homology reduction removing the majority of the data. Still, GraphPart retains more than 95% of all sequences while achieving perfect separation. Furthermore, for DNA datasets we find that a threshold of 80% maximum identity, the minimum that is achievable with CD-HIT-EST, is trivially achievable by all baseline algorithms without loss of sequences.

**Figure 3.**
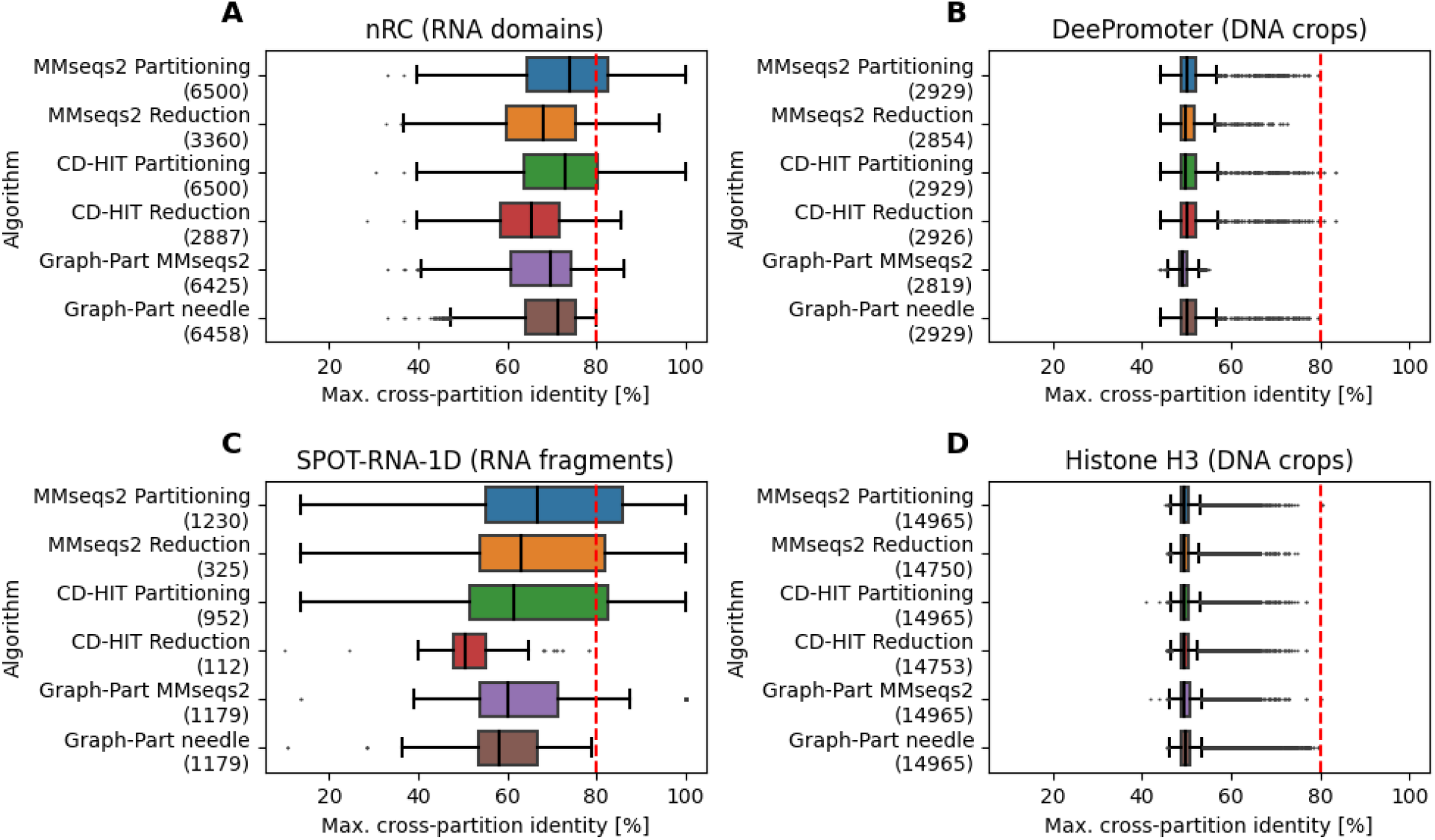
Benchmarking results on nucleotide datasets. Datasets were partitioned into five folds. The target pairwise identity threshold is indicated as the red dashed line. All algorithms were run for the generation of five equal-sized balanced partitions.

**Figure 4.**
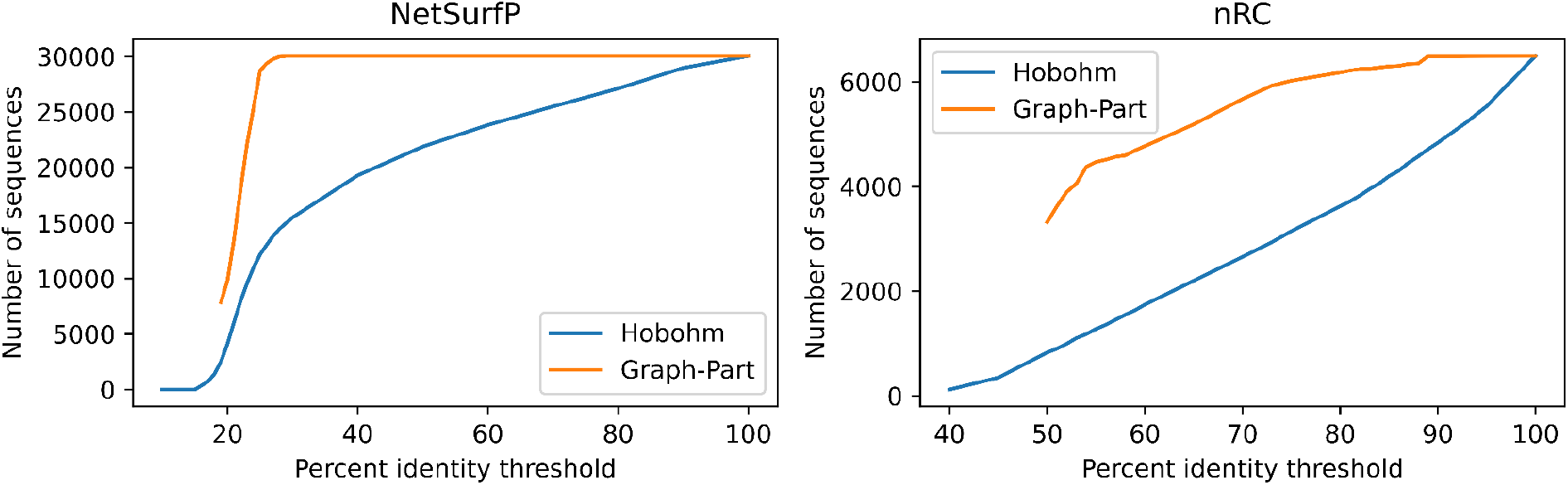
The number of sequences retained at a given maximum percent pairwise identity threshold for GraphPart and the Hobohm 2 algorithm. Needleman-Wunsch alignments were used for both algorithms, GraphPart was run for five partitions. At very low identities, GraphPart fails as heuristic iterative removal leads to the deletion of complete partitions.

While GraphPart inherently suffers from quadratic complexity due to computing a full all-vs-all similarity matrix (Supplementary Figure 1), most machine learning datasets fall within a range of size and sequence length in which application of Needleman-Wunsch alignments is feasible when using multiple cores in parallel. Furthermore, using GraphPart with MMseqs2 alignments provides a significant speedup while still comparing favorably to the baseline reduction methods.

## Conclusion

We introduce GraphPart, the first dedicated sequence homology partitioning algorithm addressing the need of homology separation in machine learning on biological sequence data. GraphPart shows superior performance over homology reduction algorithms by preserving more sequences in its partitions. Additionally, it allows for balancing of partitions for additional criteria while achieving homology separation. GraphPart is the only partitioning algorithm that guarantees its chosen identity threshold by utilizing global Needleman-Wunsch alignments. Additionally, other alignment programs such as MMseqs2 can be used by GraphPart and show competitive performance while greatly improving speed.

## Availability

GraphPart can be installed as a Python package from https://pypi.org/project/graph-part/. An online version is available at https://ku.biolib.com/graphpart. The source code is available on GitHub at https://github.com/graph-part/graph-part. All datasets and the baseline scripts used for benchmarking are included in the repository.

