## Supplementary Information for "GraphPart: Homology partitioning for biological sequence analysis"

```
Inputs: S, D, threshold
i = 1
while c>0:
      connected = []
      for s in S:
            c_other = count_cross_partition_connections(s, S, D, threshold)
            c_own = count_own_partition_connections(s, S, D, threshold)
            if c_other > c_own:
                  S = move_to_closer_partition(s,S)
            c_other = count_cross_partition_connections(s, S, D, threshold)
            if c_other > 0:
                  connected.append(s)

      c = length(connected)
      n_to_remove = c * log10(i)/100 + 1
      S, D = remove_entities(S, D, connected, n_to_remove)
      i = i + 1
```

**Code 1.** Pseudocode of the iterative removal procedure. `S` is a structure containing all entity identifiers and their associated partition membership labels. `D` contains the distances between all elements of `S`. `threshold` is a float value.

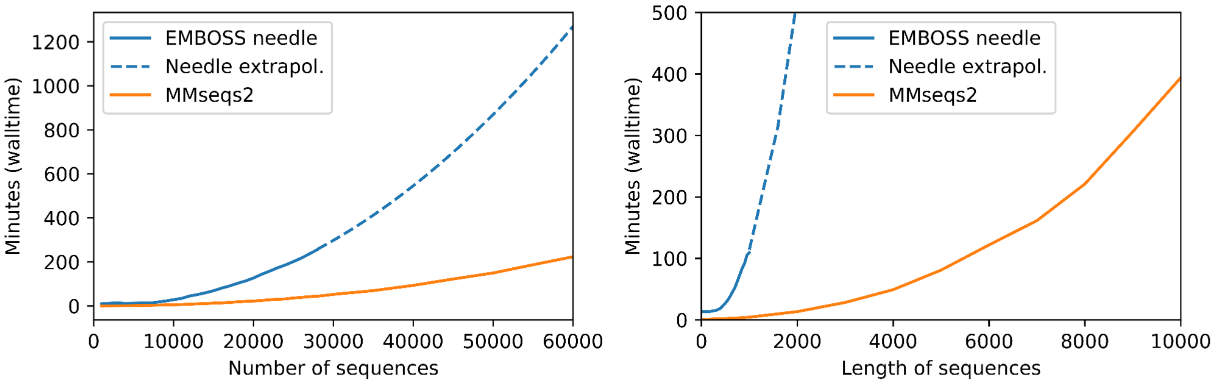


**Supplementary Figure 1.** Time complexity of GraphPart. Due to the exhaustive computation of pairwise alignments, GraphPart scales quadratically with the number of input sequences. With regards to sequence lengths m and n, pairwise alignment algorithms have a time complexity of O(mn). For the investigated dataset sizes, the execution time of the actual partitioning algorithm itself is negligible compared to the time spent on pairwise alignments.
